# Genomic landscapes of canine splenic angiosarcoma (hemangiosarcoma) contain extensive heterogeneity within and between patients

**DOI:** 10.1101/2020.11.15.380048

**Authors:** Shukmei Wong, EJ Ehrhart, Samuel Stewart, Victoria Zismann, Jacob Cawley, Rebecca Halperin, Natalia Briones, Keith Richter, Karthigayini Sivaprakasam, Nieves Perdigones, Tania Contente-Cuomo, Salvatore Facista, Jeffrey M. Trent, Muhammed Murtaza, Chand Khanna, William P. D. Hendricks

## Abstract

Cancer genomic heterogeneity presents significant challenges for understanding oncogenic processes and for cancer’s clinical management. Variation in driver mutation frequency between patients with the same tumor type as well as within individual patients’ cancers can limit the power of mutations to serve as diagnostic, prognostic, and predictive biomarkers. We have characterized genomic heterogeneity between and within patients in canine splenic hemangiosarcoma (HSA), a common naturally occurring cancer in pet dogs that is similar to human angiosarcoma (AS). HSA is a clinically, physiologically, and genomically complex canine cancer that may serve as a valuable model for understanding the origin and clinical impact of cancer heterogeneity. We conducted a prospective collection of 52 splenic masses from 44 dogs (28 HSA, 15 benign masses, and 1 stromal sarcoma) presenting to emergency care with hemoperitoneum secondary to a ruptured splenic mass. Multi-platform genomic analysis included matched tumor/normal cancer gene panel and exome sequencing. We found candidate somatic cancer driver mutations in 14/28 (50%) HSAs. Among recurrent candidate driver mutations, *TP53* was most commonly mutated (29%) followed by *PIK3CA* (14%), *AKT1* (11%), and *CDKN2AIP* (11%). We also identified significant intratumoral genomic heterogeneity, consistent with a branched evolution model, through multi-region exome sequencing of three distinct tumor regions from selected primary splenic tumors. These data provide new perspective on the genomic landscape and comparative value of understanding HSA in pet dogs, particularly as a naturally occurring cancer bearing intratumoral heterogeneity.

## Introduction

Angiosarcoma (AS), an aggressive cancer arising from vascular endothelium in anatomic sites including skin and viscera, is rare in humans, but far more common in pet dogs. Tens of thousands of canine diagnoses occur annually, comprising 45-51% of splenic cancers^1^. AS in dogs, also known as hemangiosarcoma (HSA), is a complex disease that shares clinical, histopathologic and molecular characteristics with AS^2–9^. Shared molecular features include transcriptional subtypes (angiogenic, inflammatory, and adipogenic^4,6,8^), point mutations *(TP53, PIK3CA, PTEN, PIK3R1*^8–10^), copy number gains *(PDGFRA, VEGFA, KIT, KDR),* and copy number deletions *(CDKN2A/B, PTEN*)^8,11,12^, many of which hold potential as biomarkers to guide clinical management. As a naturally occurring cancer, HSA is a setting in which to perform cross-species comparative studies to improve understanding of AS development and clinical management. AS in humans and dogs is also heterogeneous in clinical presentation, clinical course, histopathology and cellular composition. Genomic study of HSA may thereby present a unique opportunity to dissect the role of ITH in an aggressive sarcoma in which genomic biomarkers are increasingly utilized for clinical management.

Dramatic variation in clinical, cellular, and genomic cancer features is common across patients with the same tumor type as well as within tumors in individual patients. Such heterogeneity presents challenges for uniform diagnosis, prognosis, and treatment of many cancers^13^. ITH arising from branched evolution is particularly problematic for molecular and genomic diagnostics because it can lead to spatial variation in the abundance of mutations that may serve as biomarkers. Thus, especially in tumors that also bear cellular heterogeneity (e.g. with abundant stromal or vascular components), two different biopsies or even two different sections from the same biopsy may bear a variable spectrum of detectable driver mutations^14^. For example, at least 3 distinct tumor regions are needed to detect 5 key driver mutations with a 90% level of certainty^15^. ITH has also been associated with more aggressive tumor biology and poorer patient outcomes^16^. Understanding of evolutionary processes, clinical impacts, and clinical interventions in the setting of ITH is needed, but few preclinical *in vivo* models are capable of modeling such heterogeneity and human cancer studies are costly and challenging. HSA may provide a unique setting in which to study the origins and effects of ITH and its relevance for genomic diagnostics. Here, we describe the results of multi-platform genomic analysis of 51 splenic masses from 43 dogs (27 HSAs, 15 benign lesions, and 1 stromal sarcoma) including matched tumor/normal exome and cancer gene panel-based next generation sequencing. These data expand understanding of HSA’s inter- and intra-tumoral genomic heterogeneity.

## Results and Discussion

To deepen understanding of angiosarcoma’s genomic underpinnings and inter- and intra-tumoral heterogeneity, we undertook a prospective multicenter study^17^ in which genomic analysis was performed on untreated benign and malignant tissues from 43 pet dogs presenting to emergency hospitals with hemoperitoneum after splenic mass rupture. Samples and sequencing platforms are summarized in **Table 1** and **Table S1**. Of 43 cases, 27 (63%) were diagnosed with HSA, 15 (34%) were diagnosed with benign lesions (11 benign complex nodular hyperplasia, 2 complex hyperplasia with hematoma, 1 hematoma, and 1 myelolipoma) and 1 dog was diagnosed with stromal sarcoma. Tumors were collected from three geographically distinct splenic regions. Each of 3 sections was sub-divided for histopathology or genomics. Tumor content estimated via histopathology in 23/27 HSA patients ranged from 0 to > 95% (median 10%, **Table S1**). The range of tumor cell content and the high frequency of tumor sub-sections without identifiable tumor cells both underscore the degree of cellular heterogeneity in HSA and potential challenges associated with genomic discovery and diagnostic studies in this tumor type.

**Table 1.** Samples and Sequencing Platforms

| Patient Diagnosis | Total Unique Patients | Total Unique Samples | Samples per Sequencing Platforms |
| --- | --- | --- | --- |
| Hemangiosarcoma | 27 | 36* | 24 T/N CCAP, 3 T CCAP, 4 T/N exome |
| Benign Complex Nodular Hyperplasia | 11 | 11 | 3 T/N CCAP |
| Complex Hyperplasia w/Hematoma | 2 | 2 | 2 T/N CCAP |
| Hematoma | 1 | 1 | 1 T/N CCAP |
| Myelolipoma | 1 | 1 | 1 T/N CCAP |
| Stromal sarcoma | 1 | 1 | 1 T/N CCAP |
| **Total** | **43** | **52** | **33 T/N CCAP, 3 T CCAP, 4 T/N exome** |

\*Pt 3, 11, 16, 22: All 3 tumor sections exome sequenced. Pt 3: All 3 tumor sections panel sequenced. Canine patients with benign or malignant splenic mass (T) and matching normal (N) when available were sequenced on with either the Canine Cancer Amplicon Panel (CCAP) or whole exome sequenced (exome). All malignant samples were confirmed to be hemangiosarcoma except for one, a stromal sarcoma.

To identify somatic single nucleotide variants (SNVs), we utilized a custom canine cancer next generation sequencing amplicon panel^18^ with regions covering commonly mutated genes in AS and HSA (**Table S2**)^11,12,19^ including 330 exonic regions of canine orthologs of 67 commonly mutated human cancer genes. The section with the highest tumor content out of the three sections was sequenced. Matched normal and tumor tissue was sequenced to an average depth of 1,836x and 1,797x, respectively (**Table S3**). We identified 184 putative somatic SNVs across this cohort, of which 38 occurred at an allele frequency (AF) ≥ 10%. At least one somatic mutation with an AF ≥ 10% was seen in 13/27 (48%) of the sequenced tumors (**Figure 1**). The median AF for all 184 putative somatic SNVs was 0.05 (range 0.02 – 0.41). *TP53* was the most frequently mutated gene, with 11 mutations occurring across 8 patients (29%). Of 11 *TP53* mutations, 8 were missense with most occurring in human-equivalent pathogenic hotspots in the DNA-binding domain. Single cases of splice acceptor (c.530-2A>C), stop gain (R296*), and frameshift (T243fs) mutations were identified. *PIK3CA* mutations were identified in 4 patients (14%). *PIK3CA* H1047L was the only recurrent point mutation, identified in 2 cases. The third and fourth cases bore H1047R and G1049R mutations. Amino acid 1047 is the most frequently mutated *PIK3CA* hotspot in human cancers, previously shown to be mutated in HSA (30-46% of cases)^9,12,20^. *AKT1* was mutated in 3 cases (11%) with two potentially pathogenic missense mutations, G37D and R23W, and one frameshift, L52fs, with unknown significance. Additional likely pathogenic mutations in single patients included *CDKN2B* R105Q and *NRAS* Q61R. Variants of unknown significance (VUS) include *CDKN2AIP* Q67R (a gene deleted in HSA via aCGH studies^7^), an *ERBB2* splice region variant, *FLT3* N703S, *JAK3* M724T, *PTEN* H39fs, and *PTPRB* L1284P and S1965P. Of the 7 panel-sequenced benign cases, only Patient 37 (benign hyperplasia) bore a somatic SNV in a cancer gene - a VUS impacting *PTPRB.* Patient 43 (stromal sarcoma) bore a likely pathogenic missense mutation at R789C in *GNAS.* These driver mutations and their frequencies resemble those identified in HSA in other studies^9,12,20^. The genomic landscape of HSA confirmed here provides a framework to guide study of HSA development while also guiding therapeutic strategies under a precision medicine paradigm in which drug-biomarker relationships may exist in HSA (**Figure S1**). Cross-study variation in this landscape, such as the lower rate of *PIK3CA* mutations identified in our cohort relative to others, likely reflects natural variation by HSA anatomic site, clinical characteristics, and breed in addition to variation in sequencing platforms and analysis approaches. It underscores the need for expanded genomic study in very large cohorts. It also remains possible that other drivers are present in this cohort in regions not included in the panel in addition to copy number variation, translocations, and fusions. Finally, the low median AF of SNVs, despite sequencing of high tumor content sections, supports existence of significant subclonal heterogeneity.

**Figure 1.**
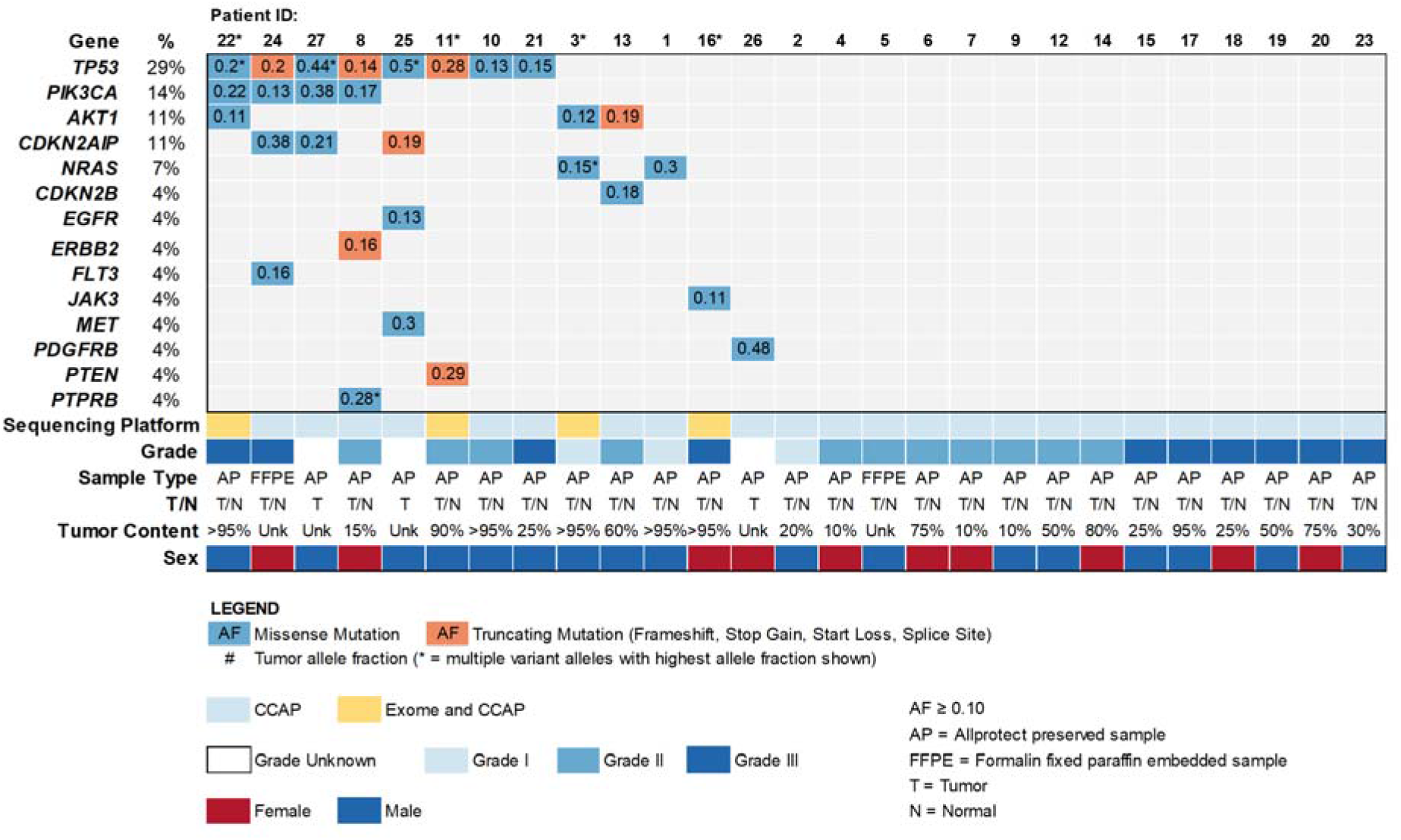
The Landscape of Somatic SNVs in Splenic Hemangiosarcoma. Recurrent, potentially pathogenic mutations were identified in 24 matched normal/tumor and 3 tumor-only FFPE- or AllProtect-preserved splenic hemangiosarcomas. As indicated by colored boxes, missense and truncating mutations identified by amplicon panel-based sequencing (CCAP) with allele frequencies ≥ 0.1 are shown. Allele frequencies are indicated in colored boxes. Tumor content percentage and tumor grade are shown where available. Patient IDs with asterisks indicate samples that were also exome-sequenced.

In order to more deeply explore heterogeneity, we next directly measured ITH via whole exome sequencing (WES) of three geographically distinct splenic sections from 4 patients (Patients 3, 11, 16, 22) along with their matching constitutional DNA from peripheral blood, achieving average sequencing coverage of 214x for tumors and 211x for normals. Patients with pathogenic mutations according to panel-based sequencing and those with high tumor content according to pathology for all 3 sections were selected (**Table S1**). These tumors bore a low tumor mutation burden (TMB), median = 0.7 mutations/Mb (range: 0.3-2.9 mutations/Mb, **Table 2**). Tumor content estimates, determined by Sequenza using AFs from exome sequencing data, suggested a median of 21% (range 10-57%, **Table 2**) whereas pathology estimates suggested a median of 75% (range 25%->95%, **Table S1**). This discordance is likely influenced by low TMB in these cases.

**Table 2.** Tumor Mutation Burden and Tumor Content of Exome-Sequenced Samples

| Sample | Tumor Section | TMB (Mutations/Mb) | Tumor Content (%) |
| --- | --- | --- | --- |
| Pt 3 | 1 | 0.315 | 10% |
| Pt 3 | 2 | 0.265 | 10% |
| Pt 3 | 3 | 1.106 | 10% |
| Pt 11 | 1 | 2.855 | 30% |
| Pt 11 | 2 | 2.582 | 52% |
| Pt 11 | 3 | 2.620 | 43% |
| Pt 16 | 1 | 0.047 | 10% |
| Pt 16 | 2 | 0.682 | 57% |
| Pt 16 | 3 | 0.702 | 40% |
| Pt 22 | 1 | 0.246 | 12% |
| Pt 22 | 2 | 1.028 | 40% |
| Pt 22 | 3 | 0.033 | 10% |

Tumor mutation burden (TMB) and tumor content of corresponding tumor sections of exome sequenced HSA patients are shown.

Analysis of intersecting somatic mutations across 3 tumor regions from 4 patients by WES identified substantial intratumoral genomic heterogeneity (**Table S6**). Between 0 and 64% of SNVs were detected in at least 2 regions and 0-13% were detected in all 3 regions. In Patient 3, 9/62 (15%) SNVs were shared in 2 or more regions and 3 (5%) were shared in all regions. *NRAS* Q61R was detected in all regions. In Patient 11, 61/181 (34%) SNVs occurred in 2 or more regions and 24 (13%) were shared among all regions. Two driver mutations, *TP53* T243fs and *PIK3CA* G1007R, were present in all regions. In Patient 16, 18/28 (64%) SNVs were shared in regions 2 and 3, including the pathogenic *TP53* Y152S. No mutations were shared by all regions or between regions 1 and 3. For Patient 22, no mutations were shared among all regions. Only region 2 bore candidate driver missense mutations (also identified by CCAP): *TP53* G256E and R272H and *PIK3CA* H1047L. In most cases where a tumor section was sequenced by both CCAP and WES, mutation concordance between CCAP-targeted and WES-targeted genomic regions was seen with the exception of *PIK3CA* G1007R in Patient 11 (detected by WES, but not CCAP likely due to its occurrence in a CCAP primer region) and AKT1 G37D in Patient 22 (detected in CCAP but not WES likely due to the increased sensitivity of CCAP sequencing). This assessment of shared or private mutations across different geographic regions from the same tumor supports that significant genomic heterogeneity exists in HSA.

To quantify the degree and makeup of ITH, we next inferred the composition of clonal groups across tumor regions using LumosVar2^21^, a tool that utilizes copy number states and point mutation allele frequencies to define clonal groups in multiple biopsies from the same patient. Each grouping is specific to the individual patient, though groups are shared among tumors from the same patient. Patient 11’s tumor bore the highest TMB and also displayed the greatest degree of heterogeneity (6 clonal groups) whereas the other 3 tumors bore 3-4 groups (**Figure 2**). Each tumor contained one unique, dominant group, Group 1, present at high frequency in all regions (40-95%). Thus, at least 40% of the cells across all tumor regions in each patient contained variants from this dominant group (**Figure 2A-D**). Those variants include all somatic mutations (including synonymous, intronic, missense, and truncating mutations). Based on their frequency, these dominant groups may be truncal or early evolutionary events in each tumor’s development. Notably, established oncogenic driver mutations such as *NRAS* Q61R (Patient 3) and *TP53* mutations (Patients 11 and 22) were typically not part of the dominant Group 1. The only Group 1 driver was *PIK3CA* G1007R in Patient 11, present in all 3 sections. In fact, no coding SNVs were present in Group 1 in Patients 3, 16, or 22 while Patient 11 contained only *PIK3CA* and *SLC10A3* (likely a passenger) variants in Group 1. Our analysis of intersecting somatic mutations from distinct regions of primary splenic HSA thus supports the presence of significant ITH consistent with a model of branched evolution in which key driver mutations may serve as early events in tumorigenesis, but these events are variable between patients. Alongside cellular heterogeneity (i.e. tumor content variability), this genomic heterogeneity holds implications for HSA development. It also presents technical challenges in mutation detection for clinical applications. Importantly, our studies focused on cases containing known drivers and modest mutation burden. Other cases with low TMB and/or lacking established drivers may well reveal different evolutionary trajectories.

**Figure 2.**
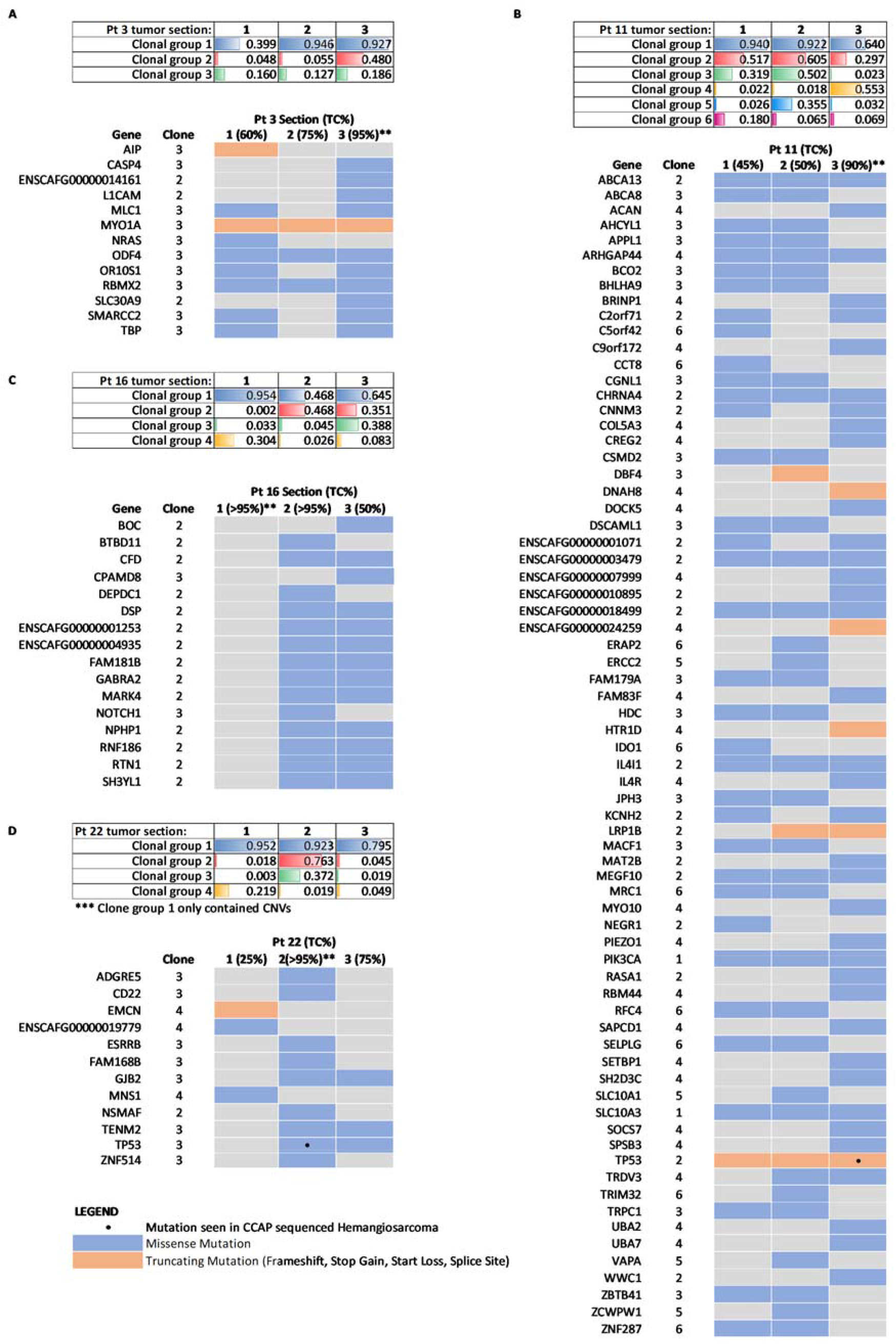
Intratumoral Genomic Heterogeneity in Splenic Hemangiosarcoma. Intratumoral heterogeneity was assessed via whole exome sequencing of 3 distinct geographic regions from 4 canine HSA patients. The composition of clonal groups across tumor regions was inferred via LumosVar2^21^ using CNVs and SNV allele frequencies. Tables show percentage of cells within each separate region for each patient within specified clonal groups. Individual patient oncoprints show only coding and nonsynonymous mutations in genes associated with each clonal group. Patients 3 (A), 11 (B) and 22 (D) have at least one missense or truncating mutation shared among all three tumor sections. Patient 16’s (C) tumor section one did not contain any SNVs that are seen in sections 2 and 3. *Total # of filtered somatic SNVs (includes all somatic SNVS: synonymous, missense, etc.). **Indicates tumor section also sequenced by amplicon panel.

ITH reflects the evolutionary trajectory of solid tumors. It may be associated with aggressive tumor biology and it presents significant challenges for genomic research and diagnostics. We have established that canine HSA contains substantial ITH. Our multi-platform genomic analysis of 52 splenic tumor samples confirms the presence of key driver mutations in HSA while also revealing extensive ITH. These data have bearing on biologic understanding and clinical management of HSA in pet dogs including shaping the development of new diagnostic tools and biomarker-driven treatment studies. These studies can also further inform the intersections in canine and human angiosarcoma genomics and present a common cancer type for studying ITH. We can now begin to leverage our understanding of the presence of this genomic heterogeneity in these naturally occurring cancer models to develop and adapt strategies to translate the value of genomic medicine to human and canine cancer patients.

## Materials and Methods

### Sample collection and nucleic acid extraction

Three splenic masses preserved in AllProtect reagent (Qiagen) and formalin from previously diagnosed HSA were used as pilot samples for evaluation of an in-house developed canine amplicon panel. For clinical samples from the 10 Ethos Discovery affiliated animal hospitals, with owner consent, whole blood and splenic masses were collected via splenectomy from 43 canine patients who were diagnosed with a hemoperitoneum with a ruptured splenic mass and had not received previous chemotherapeutic treatment for hemangiosarcoma. Tumor sample collection was performed within 5 minutes of the spleen being removed from patient. Multiple 1 x 1 x 1 cm biopsies were taken from three geographically distinct regions of the spleen (primary tumor, tumor periphery, and normal spleen). The primary tumor section was further divided into additional sections where each subsequent section was placed in the following: 10% formalin for histopathology, AllProtect reagent for nucleic acid extraction, cryovials containing fetal bovine serum and 10% DMSO for cryogenic cell storage, and L15 tissue collection media for future cell line development. The samples in formalin, AllProtect reagent, and L15 media were refrigerated at 4°C and cryovials were frozen at −20°C until submission.

2 x 10 ml of whole blood was collected into two Cell Free DNA BCT Streck tubes (Streck, Inc.) from each dog at presentation and every 60 days. Streck tubes were process as per manufacturer protocol. Buffy coat and plasma were collected and stored in −20°C until nucleic acid isolation whereas cell free DNA is stored for further investigation. Genomic DNA from 200uL buffy coat was isolated with DNeasy Blood and Tissue kit (Qiagen) according to manufacturer’s protocol. Tumor nucleic acid extractions were performed using the Allprep DNA/RNA/miRNA Universal kit (Qiagen) according to manufacturer’s protocol. Briefly, tumor tissue in Allprotect was first rinsed with sterile 1X phosphate buffered saline and minced with sterile scalpel. 30 mg of the minced tissue was then homogenized using the Bullet Blender Bead lysis kit (NextAdvance) and the resulting supernatant was further homogenized with QiaShredder (Qiagen). The flow through was used for nucleic acid extraction. Quality and quantity of blood and tumor genomic DNA was performed using the Qubit Fluorometer 2.0 (ThermoFisher Scientific) and TapeStation genomic DNA assay (Agilent Technologies). Genomic DNA was stored in −20°C until sequencing library construction. RNA was extracted and stored in −80°C for future use as per manufacturer’s instructions.

### Exome Sequencing and Analysis

Genomic DNA from blood and all 3 tumor sections from 4 HSA patients (patients 03, 11, 16, 22) underwent whole exome sequencing using a custom Agilent SureSelect canine exome capture kit with 982,789 probes covering 19,459 genes. Exome libraries were sequenced on the Illumina NovaSeq 6000 producing paired end reads of 100bp. Analysis tools and their parameters are shown in **Table S4**. FASTQ files were aligned to the canine genome (CanFam3.1.75) using BWA v0.7.8. Aligned BAM files were realigned and recalibrated using GATK v3.3.0 and duplicate pairs were marked with Picard v1.128 (http://broadinstitute.github.io/picard). Somatic single nucleotide variants (SNV) were identified only when called by two or more of the following callers: Seurat v2.6, Strelka v1.0.13 and MuTect v1.1.4. Germline SNVs were called using Haplotype Caller (GATK v3.3.0), Freebayes and samtools-Mpileup. Variant annotation was performed with SnpEff v4.3. TMB was calculated as the total number of somatic mutations per haploid callable Megabase (Mb) from WES^22^. Percent tumor content was inferred from somatic mutation data utilizing Sequenza.^23^ LumosVar2^21^ was utilized for clonal variant group analysis. Clonal groups may contain both CNVs and SNVs. Only SNVs designated as “PASS SomaticDetected” were utilized for clonal analysis. The exome data corresponding to normal blood and tumor sections are uploaded to Sequencing Read Archive under BioProject PRJNA677995.

### Targeted Amplicon Sequencing and Analysis

Targeted amplicon sequencing was performed on genomic DNA from matched blood and DNA from individual patient tumor section with the highest tumor content. A custom canine HSA cancer amplicon sequencing panel consisting of 330 amplicons targeting exons and hotspot regions in 68 genes, with amplicon sizes ranging from 91-272 bp was developed (**Table S2**). Primers were pooled in two multiplexed pools to separate adjacent amplicons and any amplicons with high potential for cross-amplification using *in silico* PCR. Sequencing libraries were constructed using droplet-based PCR amplification following the manufacturer’s protocols for the ThunderBolts Cancer Panel with specific modifications (RainDance Technologies) as previously described^24^. Paired-end sequencing was performed on an Illumina MiSeq generating 275bp reads. Analysis tools and parameters are shown in **Table S4**. Sequencing reads were demultiplexed and extracted using Picard 2.10.3. Sequencing adapters were trimmed using ExpressionAnalysis ea-utils and quality of fastq files were assessed with FastQC v0.11.5. Sequencing reads were aligned to CanFam3.1.75 using bwamem-MEM. Custom in-house scripts based on SAMtools were used to create pileups for every sample. Pileups were analyzed in R to call SNVs and indels. For each potential non-reference allele at each targeted locus in a sample, we evaluated the distribution of background noise across all other sequenced samples. To call a variant, we required the observed non-reference allele is an outlier from the background distribution with a Z-score > 5. In addition, we required tumor depth ≥100x, allele frequency ≥10%, number of reads supporting the variation ≥10, and allele fraction in the germline sample <1%. Finally, variant calls were manually curated by visualization in IGV v2.4.9. All sequencing data have been deposited in the Sequencing Read Archive under BioProject PRJNA677995.

## ACKNOWLEDGEMENTS

We thank the families of the pets who contributed samples for this study. This study was funded by Ethos Discovery and philanthropic contributions to the TGen Foundation.

## AUTHOR CONTRIBUTIONS

Conception and design: C. Khanna, W.P.D. Hendricks, S. Stewart, E.J. Ehrhart

Development of methodology: M. Murtaza, W. Hendricks, E.J. Ehrhart

Acquisition of data (acquired and managed patients, provided tissues, processed tissues, performed sequencing, etc.): S. Stewart, S. Wong, T. Contente-Cuomo

Analysis and interpretation of data (e.g., statistical analysis, biostatistics, computational analysis): S. Wong, R. Halperin, N. Briones, K. Sivaprakasam, N. Perdigones, S. Facista

Writing, review, and/or revision of the manuscript: S. Wong, C. Khanna, J. Cawley, W.P.D. Hendricks

Administrative, technical, or material support (i.e., reporting or organizing data, constructing databases): V. Zismann, Y. Pan, Y. He, K. Richter, J. Trent

Study supervision: C. Khanna, W.P.D. Hendricks

---

**Figure S1. The Naturally Occurring Heterogeneity of Canine Hemangiosarcoma Provides a Unique Opportunity to Test Hypotheses Addressing Challenges in the Delivery of Cancer Precision Medicine across Species**. Intratumoral genomic heterogeneity presents challenges to cancer precision medicine (i.e. the utilization of genomic diagnostics to guide the clinical management of cancer patients). For example, variability in subclonal frequency of driver mutations may modulate treatment response. Additionally, such variability means that any given biopsy utilized for genomic analysis may not contain driver mutations with predictive associations even if these mutations are highly abundant in major lineages in the tumor. This study has confirmed the presence of genomic variants and intratumoral heterogeneity in canine HSA that can facilitate hypothesis testing and methods development to meet these needs. For example, NRAS and PIK3CA mutations such as those described in HSA have been associated with MEK and PIK3CA inhibitor responses, respectively, in human cancers and may also be associated with such responses in canine cancers. Such responses, however, may also be dependent on intratumoral heterogeneity and will certainly be dependent on the ability to detect subclonal or low allele-frequency mutations. Basket and umbrella clinical trials that test such hypotheses in canine HSA will hold value broadly for understanding heterogeneous human cancers and will be especially valuable in less common human cancers such as angiosarcoma.

## SUPPLEMENTAL TABLES

**Table S1. Extended Demographic and Clinical Annotation**

**Table S2. Genes Included in Custom Cancer Amplicon Panel**

**Table S3. Targeted Panel and Exome Sequencing Statistics**

**Table S4. Informatic Tools, Versions, and Parameters Utilized in Primary Analysis of Canine Hemangiosarcoma Whole Exome Data**

**Table S5. Somatic SNVs Identified by Panel Sequencing in Canine Splenic Lesions**

**Table S6. Somatic SNVs Identified by Exome Sequencing in HSA**

**Table S7. Intratumoral Clonal Variant Groups Determined by Analysis of Somatic Variants Identified by Whole Exome Sequencing.**

**Table S8. Somatic CNVs Identified by Exome Sequencing in Canine Hemangiosarcoma**

## REFERENCES

1. Vail, D.M., Thamm, D. & Liptak, J. Withrow and MacEwen’s Small Animal Clinical Oncology-E-Book, (Elsevier Health Sciences, 2019).

2. Vail, D.M. & MacEwen, E.G. Spontaneously occurring tumors of companion animals as models for human cancer. Cancer Invest 18, 781–92 (2000).

3. Dickerson, E. et al. Mutations of phosphatase and tensin homolog deleted from chromosome 10 in canine hemangiosarcoma. Veterinary pathology 42, 618–632 (2005).

4. Tamburini, B.A. et al. Gene expression profiling identifies inflammation and angiogenesis as distinguishing features of canine hemangiosarcoma. BMC cancer 10, 619 (2010).

5. Behjati, S. et al. Recurrent PTPRB and PLCG1 mutations in angiosarcoma. Nature genetics 46, 376–379 (2014).

6. Gorden, B.H. et al. Identification of three molecular and functional subtypes in canine hemangiosarcoma through gene expression profiling and progenitor cell characterization. Am J Pathol 184, 985–995 (2014).

7. Thomas, R. et al. Genomic profiling reveals extensive heterogeneity in somatic DNA copy number aberrations of canine hemangiosarcoma. Chromosome research 22, 305–319 (2014).

8. Megquier, K. et al. Comparative genomics reveals shared mutational landscape in canine hemangiosarcoma and human angiosarcoma. Molecular Cancer Research 17, 2410–2421 (2019).

9. Wang, G. et al. Molecular subtypes in canine hemangiosarcoma reveal similarities with human angiosarcoma. PloS one 15, e0229728 (2020).

10. Wang, G. et al. Actionable mutations in canine hemangiosarcoma. PloS one 12 (2017).

11. Thomas, R. et al. Genomic profiling reveals extensive heterogeneity in somatic DNA copy number aberrations of canine hemangiosarcoma. Chromosome Res 22, 305–19 (2014).

12. Wang, G. et al. Actionable mutations in canine hemangiosarcoma. PLoS One 12, e0188667 (2017).

13. Melo, F.D.S.E., Vermeulen, L., Fessler, E. & Medema, J.P. Cancer heterogeneity—a multifaceted view. EMBO reports 14, 686–695 (2013).

14. Dagogo-Jack, I. & Shaw, A.T. Tumour heterogeneity and resistance to cancer therapies. Nature reviews Clinical oncology 15, 81 (2018).

15. Sankin, A. et al. The impact of genetic heterogeneity on biomarker development in kidney cancer assessed by multiregional sampling. Cancer medicine 3, 1485–1492 (2014).

16. Morris, L.G. et al. Pan-cancer analysis of intratumor heterogeneity as a prognostic determinant of survival. Oncotarget 7, 10051 (2016).

17. Stewart, S.D., Ehrhart, E., Davies, R. & Khanna, C. Prospective observational study of dogs with splenic mass rupture suggests potentially lower risk of malignancy and more favorable perioperative outcomes. Veterinary and Comparative Oncology (2020).

18. Lorch, G. et al. Identification of frequent activating HER2 mutations in primary canine pulmonary adenocarcinoma. bioRxiv, 528182 (2019).

19. Behjati, S. et al. Recurrent PTPRB and PLCG1 mutations in angiosarcoma. Nat Genet 46, 376–379 (2014).

20. Lindblad-Toh, K. Genomic analysis reveals shared genes and pathways in human and canine angiosarcoma. bioRxiv (2019).

21. Halperin, R.F. et al. Leveraging spatial variation in tumor purity for improved somatic variant calling of archival tumor only samples. Frontiers in oncology 9, 119 (2019).

22. Xu, Z. et al. Assessment of tumor mutation burden calculation from gene panel sequencing data. Onco Targets Ther 12, 3401–3409 (2019).

23. Favero, F. et al. Sequenza: allele-specific copy number and mutation profiles from tumor sequencing data. Ann Oncol 26, 64–70 (2015).

24. Murtaza M, D.S., Pogrebniak K, Rueda OM, Provenzano E, Grant J, Chin SF, Tsui DW, Marass F, Gale D, Ali HR, Shah P, Contente-Cuomo T, Farahani H, Shumansky K, Kingsbury Z, Humphray S, Bentley D, Shah SP, Wallis M, Rosenfeld N, Caldas C. Multifocal clonal evolution characterized using circulating tumour DNA in a case of metastatic breast cancer. Nat Commun 4, 8760 (2015).

