## Supplementary figures and images for "Genomic landscapes of canine splenic angiosarcoma (hemangiosarcoma) contain extensive heterogeneity within and between patients"

### Figure S1

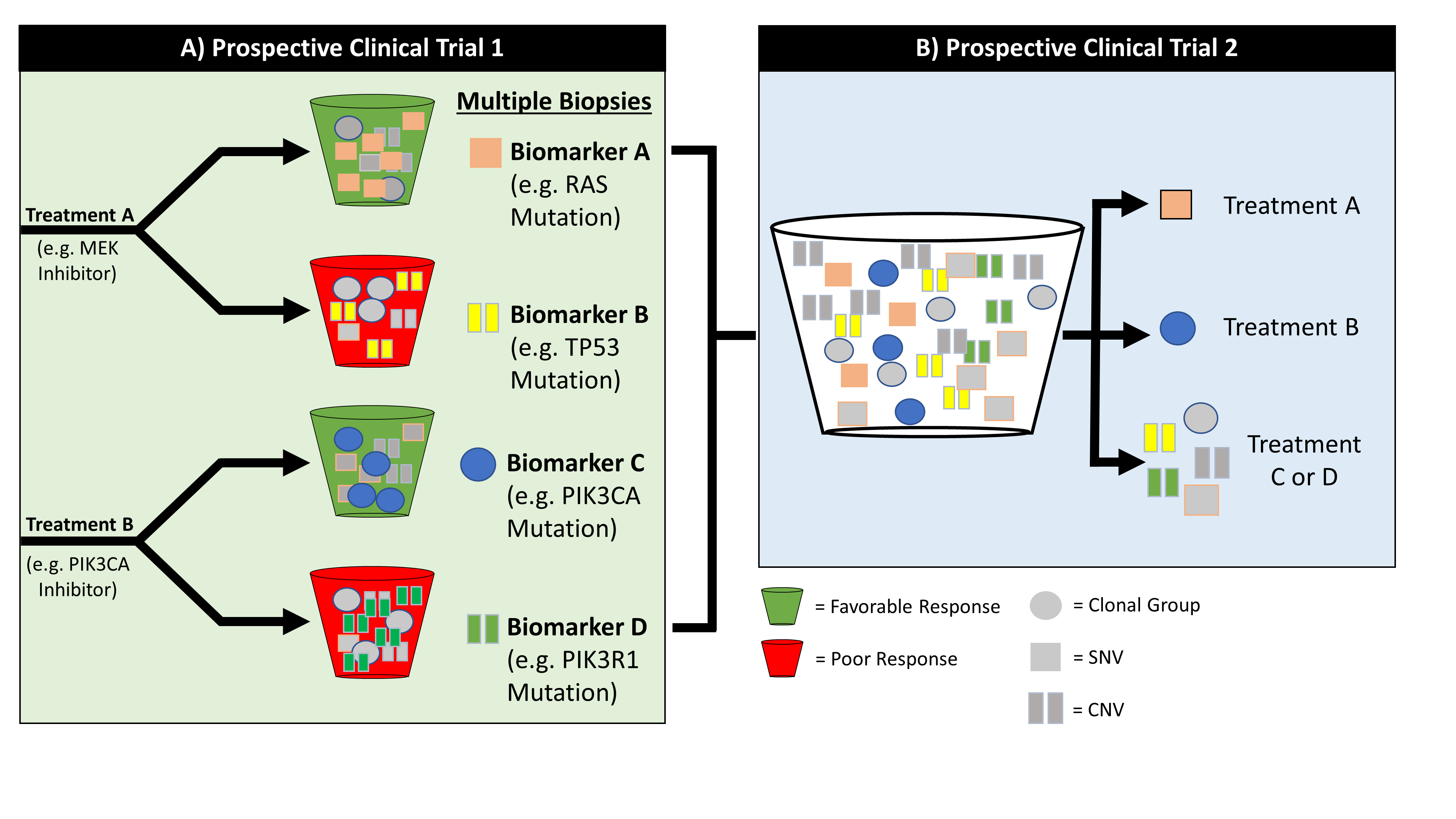
